# A multistate capture-recapture model to estimate reproduction of North Atlantic right whales

**DOI:** 10.1101/2024.04.13.589367

**Authors:** Daniel W. Linden, Richard M. Pace, Lance P. Garrison, Jeffrey A. Hostetler, Amy R. Knowlton, Véronique Lesage, Rob Williams, Michael C. Runge

## Abstract

The recent steep decline of critically endangered North Atlantic right whales (*Eubalaena glacialis*) can be attributed to high mortality combined with low reproduction. While the former is a clear result of anthropogenic activity, the latter involves more complexity. Evidence suggests that both short-term fluctuations in prey availability and long-term decline in health are responsible for depressed right whale calving rates. To facilitate an assessment of extinction risk, we developed a multistate capture-recapture model that estimated the probability of calving using extensive sightings data from 1990-2019. The model estimated sub-lethal effects of severe injury on calving probability, and modeled temporal variability in calving as related to indices of prey availability (*Calanus* spp. biomass) and an apparent regime shift. The average annual probability of calving for known-breeding females, given average prey conditions, decreased from 0.217 [0.162, 0.281] to 0.142 [0.067, 0.252] after the 2010 regime shift. The model indicated strong evidence of a relationship between calving probability and the prey index from the eastern Gulf of Maine, though this relationship effectively disappeared after 2010; moderate evidence for a relationship with prey from the southwest Gulf of St. Lawrence remained. Weak evidence of reduced calving probability due to severe injury resulted from low sample sizes, given increased mortality for individuals observed with severe injuries. The regime effect is hypothesized to be capturing a long-term decline in health due to a combination of decreasing habitat quality resulting from climate change and potentially chronic sublethal injuries (e.g., entanglements). Our reproduction model provides demographic parameter estimates that can be used in population projections for North Atlantic right whales, though uncertainty remains in the mechanisms responsible for recent declines in calving.

## 1. Introduction

The recent steep decline of critically endangered North Atlantic right whales (*Eubalaena glacialis*; hereafter, NARWs) has been a consequence of unusually high mortality rates combined with low calf production (Hayes et al., 2023). Documented mortalities of NARWs, aside from neonatal loss, are primarily due to human-caused interactions (Sharp et al., 2019; Knowlton et al., 2022), given heavy overlap between the species distribution and anthropogenic activity in the western North Atlantic Ocean (Kraus and Rolland, 2007). While the increased NARW deaths over the past decade are alarming and consequential (Davies and Brillant, 2019), the risks of vessel strike and fishing gear entanglement have long been documented in this population (Kraus, 1990; Knowlton and Kraus, 2001). The convergence of high mortality and low calving rates during the 2010s may be related to an apparent distribution shift by NARWs (Davis et al., 2017; Simard et al., 2019; Meyer-Gutbrod et al., 2023); it is unclear whether the shift represents a short-term anomaly or a permanent change for the species.

Prey availability has historically influenced both NARW distribution and calving. The primary prey for NARWs includes the late developmental stages of the copepod *Calanus finmarchicus* (Mayo and Marx, 1990), which needs to occur in dense patches to serve as an effective food source (Baumgartner et al., 2007). Decadal fluctuations in relative *C. finmarchicus* abundance have exhibited a strong relationship with NARW calving rates (Greene and Pershing, 2004; Meyer-Gutbrod et al., 2015), including positive anomalies in the 1980s and 2000s, and negative anomalies in the 1990s. These anomalies in prey abundance were linked to Arctic climate regimes and ocean circulation patterns (Greene et al., 2008). Other copepod species have been associated with NARW seasonal distribution and abundance (Pendleton et al., 2009), including the larger *C. hyperboreus*, which dominates the copepod biomass in the Gulf of St. Lawrence [GSL; Sorochan et al. (2019)] and may explain the recent importance of this area as NARW habitat (Simard et al., 2019; Crowe et al., 2021). Use of the GSL by mature females was associated with higher reproductive success during 2016–2021 (Bishop et al., 2022), though bioenergetics modeling suggests a decline in the energy provided by *Calanus* spp. biomass and overall habitat suitability in the GSL during the 2010s compared to the 2000s (Lehoux et al., 2020; Gavrilchuk et al., 2021). Changes in whale distribution in response to apparent resource availability may not be sufficient to overcome large-scale effects of climate forcing (Meyer-Gutbrod et al., 2021).

Other long-term trends in NARW health, body condition, and calving indicate significant challenges for the prospect of species recovery. Body lengths of NARWs have decreased since 1981 (Stewart et al., 2021), and smaller body sizes correspond with reduced NARW calving probability (Stewart et al., 2022; Pirotta et al., 2024). Observations of NARW body condition (Rolland et al., 2016; Pettis et al., 2017) and statistical models examining the cumulative effects of sub-lethal stressors on NARW health (Schick et al., 2013; Knowlton et al., 2022; Pirotta et al., 2023) all indicate long-term declines during the past 30+ years that are associated with increased calving intervals and decreased overall reproduction. Differences in blubber thickness between NARW and southern right whales (SRW; *Eubalaena australis*) suggests a nutritional deficiency in NARW which makes reproduction more variable and limits population growth potential (Miller et al., 2011). Calf production by NARW indicated an annual rate of increase that was 1/3 of that observed for SRW populations globally; while most of this difference was attributed to reduced adult female NARW survival (due to human-caused mortalities), projections under high survival were still unable to match the realized growth rates of SRW (Corkeron et al., 2018). The suppressed reproduction in NARW appears to be a complex result of anthropogenic stressors and climate-associated changes in the quality and availability of prey (Meyer-Gutbrod et al., 2021).

Our objective in this paper was to estimate demographic rates related to NARW calf production and its relationships with severe injury and prey availability to inform a population evaluation tool that could help facilitate recovery efforts (Moore et al., 2021). Previously, we estimated cause-specific mortality in NARWs to better understand the lethal anthropogenic threats to the population (Linden et al., 2023). Here, we developed a multistate capture-recapture model (Lebreton et al., 2009) to estimate the probability of calving as a function of age and breeding state, and also calf survival during the first ∼6 months after birth, all while accommodating uncertainty in the observation processes. We also estimated sub-lethal effects of severe injury on calving probability, and modeled temporal variability in calving as related to indices of *Calanus* spp. (Sorochan et al., 2019) and two potential regime shifts related to changes in the environment and NARW response to the changes. The collection of parameter estimates was intended to provide a baseline reference for population projections (Runge et al., 2023) that could help assess threats to NARW persistence in the context of changes in human activities and environmental conditions in the western North Atlantic Ocean.

## 2. Materials and methods

### 2.1. Monitoring data

Monitoring efforts by the North Atlantic Right Whale Consortium (https://www.narwc.org/) have covered most of the species range in the western North Atlantic since 1980 (Kraus and Rolland, 2007), though consistent effort began closer to 1990 (Pace et al., 2017). Directed aerial surveys with trained observers flown in heavily used habitats throughout the species range comprise the bulk of sightings. NARWs are individually identifiable due to distinct natural markings (i.e., callosity patterns) that become permanent at a young age (>0.5-years old); the catalog curated by the New England Aquarium includes high quality photographs that allow for constructing individual sightings histories (Hamilton et al., 2007). In addition to identifying individuals, a review of all photographs to look for evidence of human related scarring and to assess health is conducted annually (Knowlton et al., 2016; Pettis et al., 2017); the severity of observed injuries and various health metrics are scored. Data on recovered carcasses of large whales are gathered and maintained by multiple stranding networks situated along the Atlantic coasts of the United States (*Greater Atlantic Marine Mammal Stranding Network* | *NOAA Fisheries*) and Canada (*Quebec Marine Mammal Emergency Response Network* and *Marine Animal Response Society*). We relied on a detailed list of documented right whale mortalities aggregated from those networks at the Northeast Fisheries Science Center (Henry et al., 2021).

We used sightings data extracted on 23 December 2021, which included 691 whales >0.5-years old known to be alive sometime during 1 April 1990–30 September 2019. We used these same data in a different framework to estimate cause-specific rates of injury and mortality (Linden et al., 2023). In this paper, we constructed encounter histories by individual (n=691) and year (T=30) with a primary focus, instead, on the presence of a dependent calf during the summer. We defined an encounter period of 1 April–30 September (i.e., “summer” surveys), during which observations of individual states (or events) could occur. The recorded event *y*_*i,t*_ for an individual *i* in year *t* could take one of three values: 1 = seen alive; 2 = seen alive with a calf; and 3 = not seen.

In addition to the defined encounter period, we used sightings collected from the southeastern United States coast during the previous calving season (1 Dec–30 Mar; “winter” surveys) to provide further evidence of an individual’s reproductive state. The winter surveys are believed to provide a near-complete census of calf production (Kraus and Rolland, 2007) and, as such, sightings of calves allowed us to identify the true reproductive state of the associated adult females. By defining our encounter histories with regards to the summer survey period, we enabled the estimation of calf loss during the first ∼6 months (with accommodations for imperfect detection).

Other individual attributes included age class (i.e., birth year) and sex, which were known for 89% and 94% of individuals, respectively. In addition, severe injury status was assigned in the year that the injury was first detected based on photographic assessment (A. Knowlton, New England Aquarium, *unpublished data*, 2021). Severe injuries included entanglements that were classified as causing a severe wound (sometimes with gear attached), and vessel strike wounds that were deep (Pettis et al., 2017).

### 2.2. A multistate model for reproduction and survival

#### 2.2.1. State transition matrix

Our multistate capture-recapture model described transitions between individual states to estimate probabilities of survival and reproduction (Figure 1). State transition probabilities were conditional on multiple individual attributes (see Figure S1 for a representation specific to age classes) and varied over time. We defined the following true reproductive states: 1 = alive and male; 2 = alive and female, nulliparous; 3 = alive and female, with calf; 4 = alive and female, resting from recent calf event; 5 = alive and female, waiting for next calf event; and 6 = dead. All females were either nulliparous [i.e., pre-breeders; Reed et al. -Reed et al. (2022)] or at some stage in the breeding cycle, acknowledging that females must rest for at least 1 year after weaning a calf before waiting to breed and birth another calf (Hamilton et al., 1998; Kraus and Rolland, 2007).

**Figure 1.**
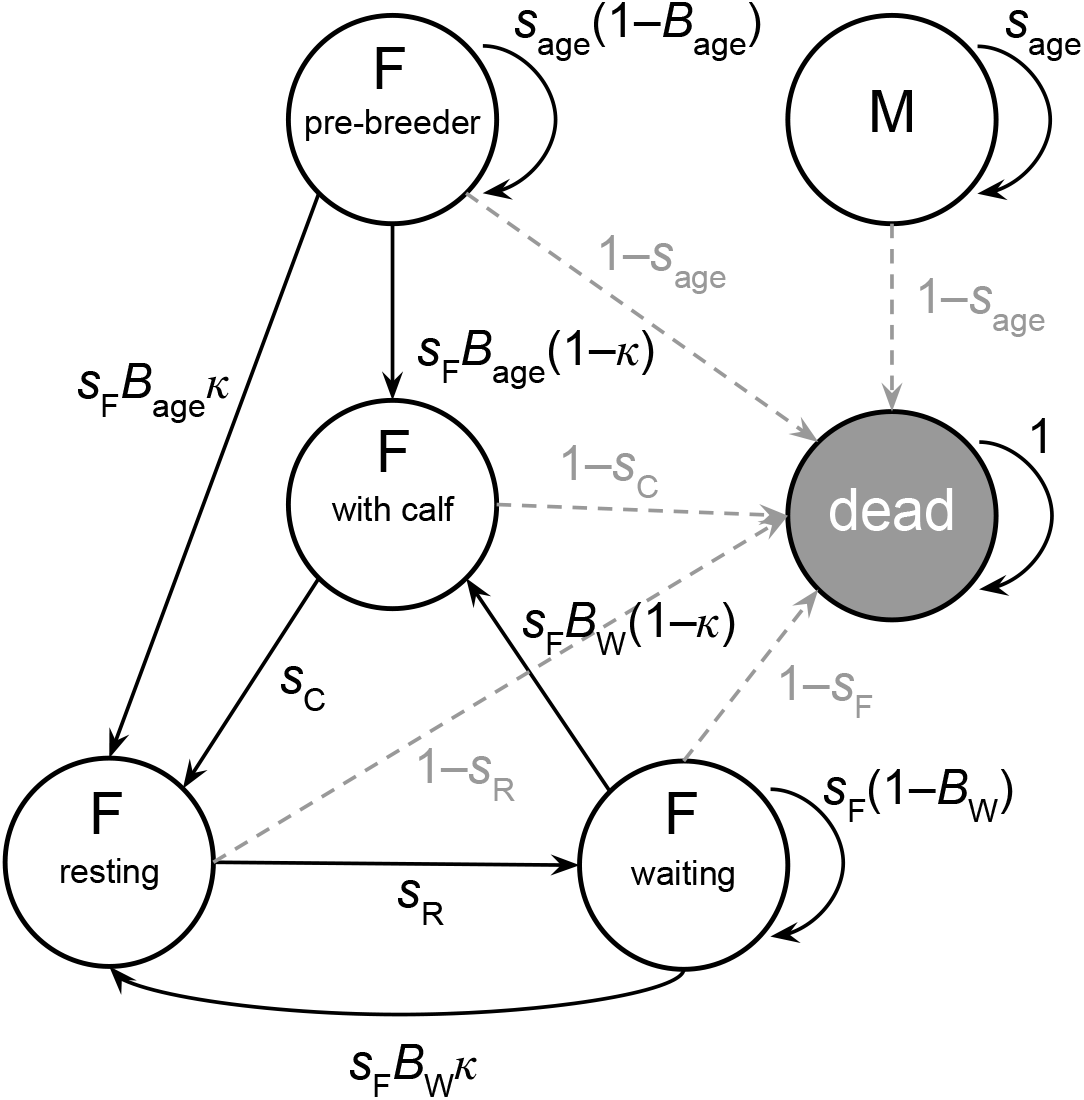
Simplified state transition model for North Atlantic right whale reproductive states across annual time steps. Transitions between states from year *t* to *t* + 1 depend on probabilities of survival (*s*) and reproduction (*B*), which are conditional on the state and age class of an individual in year *t*. Subscripts for indexing individual *i* and year *t* are omitted for clarity.

The mean age-specific survival probabilities for individuals younger than 4.5 years old were shared for males and females (e.g., *s*_1_,…,*s*_4_) until reaching an adult age when survival became sex specific (*s*_*F*_, *s*_*M*_). The age cutoff for adults was chosen based on previous NARW modeling (Pace et al., 2017) and the earliest observation of breeding in a known-age female (5 yrs). The physiological demands of breeding warranted separate mean survival probabilities for females with a calf (*s*_*C*_), for females resting from having recently weaned a calf (*s*_*R*_), and for females waiting to breed (*s*_*F*_). Along with survival probabilities, transitions between reproductive states for females were determined by mean probabilities of first breeding (*B*_*age*_) and continued breeding (*B*_*W*_), and the probability of early calf loss (*κ*). Early calf loss (death of calf within ∼6 months of birth) resulted in an individual skipping the “with calf” state and transitioning directly to the resting state.

Aside from reproductive state and age class, we used additional individual and temporal features to explain variation in survival and breeding probabilities. As such, the probabilities were defined as individual- and year-specific, resulting in the following state transition matrix:

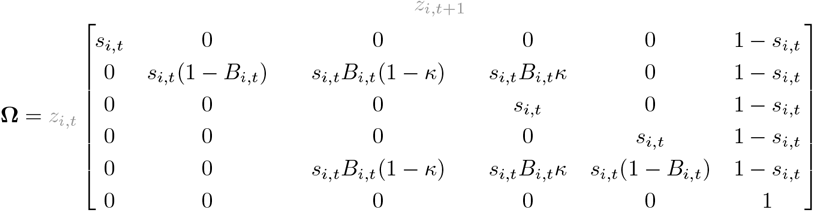

Males that survive remain in state 1, while females that survive transition between the various reproductive states (i.e., *z*_*i,t*+1_ = 2, 3, 4, or 5) according to the probabilities of reproduction (*B*_*i,t*_) and calf loss (*κ*). Death (*z*_*i,t*+1_ = 6) is an absorbing state.

#### 2.2.2. Estimating variation in probabilities of state transitions

We used generalized linear models to estimate variation in probabilities of survival and reproduction. Survival probability was modeled using a hazard rate formulation (Ergon et al., 2018) such that 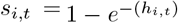, where *h*_*i,t*_ is the mortality hazard rate for individual *i* from year *t* to year *t* + 1. A log-linear model for the mortality hazard rate was specified as:

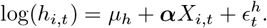

Here, *µ*_*h*_ is the log-scale average hazard rate for adult males and nulliparous females ≥ 4.5 years old; ***α*** is a vector of effects coefficients representing the log-scale differences in mortality rate for various individual and temporal attributes; *X*_*i,t*_ is a matrix of indicator variables for said attributes specific to individual *i* and year *t*; and 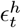 is the random normal effect of year *t* on the hazard rate. The effects included those for ages of young animals, ranging from age 1 (0.5 yr old) to age 4 (e.g., *α*_1_, …, *α*_4_); the reproductive stages of being with calf (*α*_C_) and resting from a recent calf (*α*_R_); presence of a severe injury (*α*_inj_); and a regime shift in 2013 (*α*_2013regime_). The 2013 regime shift reflected a period that had resulted in fewer sightings from traditional survey locations due to an apparent shift in the whale distribution (Davis et al., 2017; Simard et al., 2019) and, in particular, detections of NARW from a long-term monitoring program in the northern Gulf of St. Lawrence (Meyer-Gutbrod et al., 2023).

We used a logit-linear model for probability of reproduction, which was conditional on an individual being an adult female (≥ 4.5 years old):

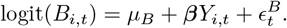

Here, *µ*_*B*_ is the logit-scale average probability of calving for females that were proven breeders (known to have had a calf before); ***β*** is a vector of effects coefficients representing the logit-scale differences in calving probability for various individual and temporal attributes; *Y*_*i,t*_ is a matrix of indicator variables for said attributes specific to individual *i* and year *t*; and 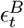 is the random normal effect of year *t*. The effects included age-specific calving for first-time breeders, ranging from age 5 to age 10+ (e.g., *β*_5_, …, *β*_10+_); presence of a severe injury (*β*_inj_); a regime shift in 2010 for all adult females (*β*_2010regime_), reflecting the major ecosystem shift in the western Atlantic; relative prey availability as indexed by two locations, the southwest Gulf of St. Lawrence (*β*_prey=GSL_) and the eastern Gulf of Maine (*β*_prey=GOM_); and finally, an interaction between the regime shift and the GOM index (*β*_prey=GOM|2010regime_).

Relative prey availability was represented by annual indices of *Calanus* spp. biomass (mg m^−2^) at various locations in the western North Atlantic (east coast waters of the United States and Canada) as calculated by Sorochan et al. (2019). Annual values of biomass for late state copepods were derived for several sub-regions in the GOM and the GSL using general linear models of spatially and seasonally variable zooplankton samples. We obtained an augmentation of the published time series that included data from 2017-2019 (K. Sorochan, Fisheries and Oceans Canada, *unpublished data*, 2022). Preliminary analyses (*see Supplementary material*) identified the southwest GSL and the eastern GOM locations as capable of explaining variation in per-capita calving when calculated as the log-transformed mean of a 3-year moving window (Figure 2). The window included the focal year and 2 years prior (i.e., *t, t* − 1, *t* − 2) and roughly coincided with an optimal 3-4-year calving interval for NARWs (Hamilton et al., 1998). We hypothesized that the *Calanus* spp. indices represented a multi-year feature of prey conditions that helped determine whether individual females would have the energy stores to reproduce. We included the interaction between the 2010 regime shift and the GOM index given the overwhelming evidence of ecosystem change in the GOM.

**Figure 2.**
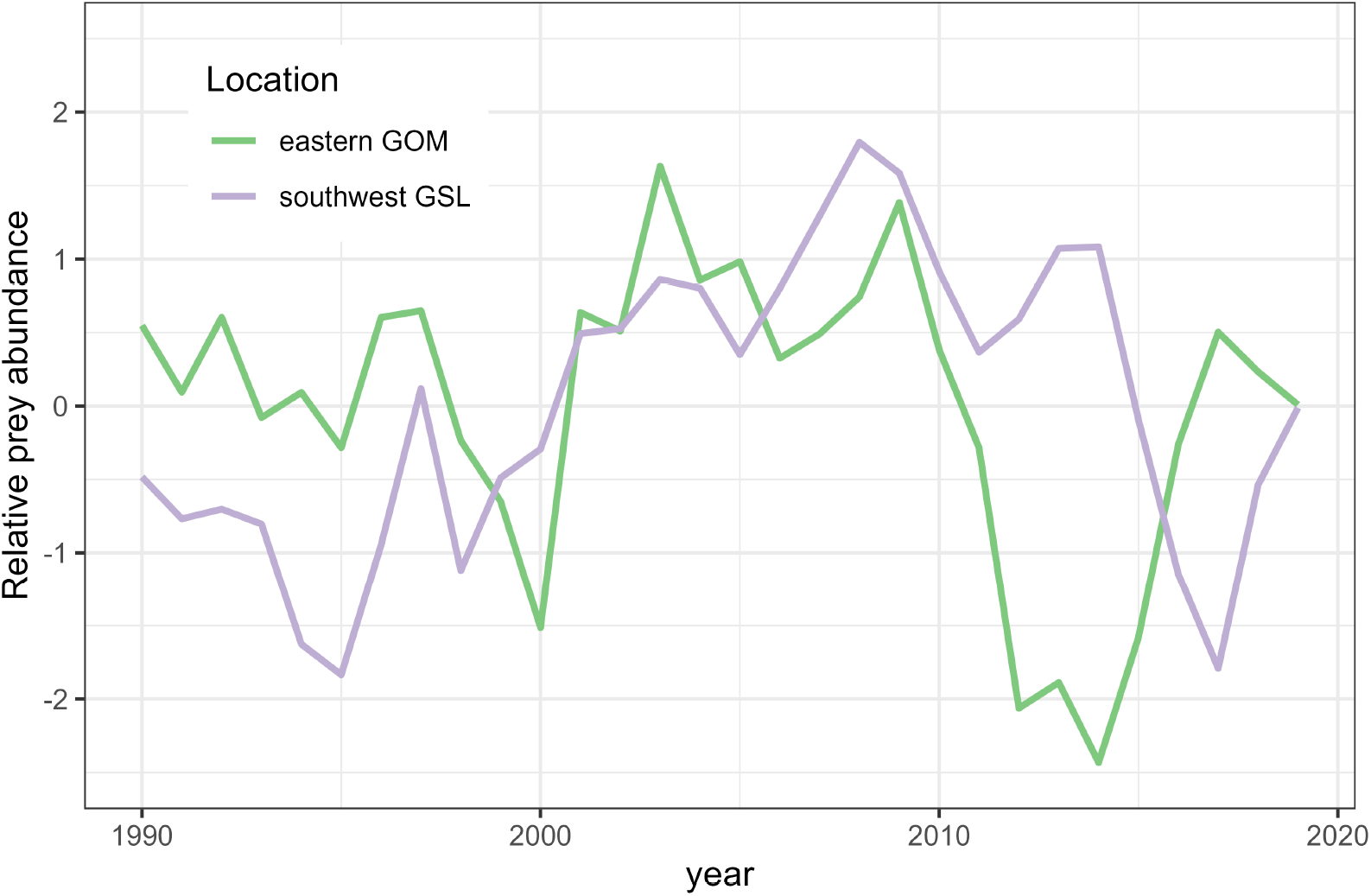
Relative prey abundance (3-yr moving average of log *Calanus* spp. biomass in mg m^2^) for the North Atlantic right whales in the southwest Gulf of St. Lawrence (GSL) and the eastern Gulf of Maine (GOM), 1990-2019.

#### 2.2.3. Observation matrix

The observations *y*_*i,t*_ were conditional on true states *z*_*i,t*_ as dictated by the observation matrix (Θ), which linked each true state to the possible observations that could occur in a given year. Sighting probabilities (*p*_*i,t*_) varied by individual and year. The observation matrix was thus defined as follows:

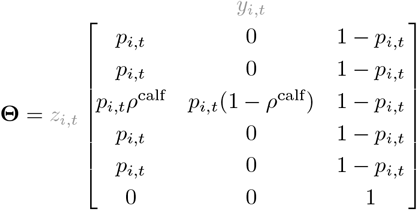

Individuals that are alive are sighted with probability *p*_*i,t*_, while *ρ*^calf^ represents the probability of missing a calf in the summer. We modeled variation in sighting probabilities using a logit-linear model:

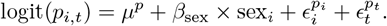

Here, *µ*^*p*^ is the logit-scale mean sighting probability; *β*_sex_ is the effect of the contrast variable sex_*i*_ (−1 = male, 1 = female); 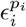 is the random normal effect of individual *i*; and 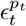 is the random normal effect of year *t*.

### 2.3. Model fitting and assessment

We fit the model in R (R Core Team, 2022) using Markov chain Monte Carlo (MCMC) with NIMBLE (de Valpine et al., 2017, 2022). We conditioned on first capture (f_*i*_) and used known breeding states as data. As an example of known breeding states, if a female was seen during the summer in year *t* with a 0.5-year-old calf and then subsequently seen alive in year *t* + 2, the known breeding states would have been *z*_*t*_ = 3, *z*_*t*+1_ = 4, and *z*_*t*+2_ = 5. Using known states as data greatly increases computational efficiency (Kéry and Schaub, 2012). This strategy also allowed us to directly incorporate carcass recoveries, where *z*_*t*_ = 6 for identified animals that were found dead. The general likelihood for the sightings data was therefore defined as follows:

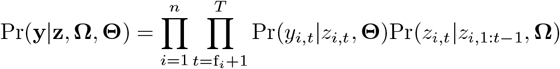

The likelihood was also conditional on individual attributes (i.e., sex, age, and injury status) and imputation was necessary for some missing values. Individuals with an unknown initial age or sex were assigned values according to probability distributions. The distribution of observed known ages was not informative for unknown ages given that known-age individuals had either been sighted as 0.5-yr-old animals or had ancillary information available (e.g., genetic sample or sighting prior to 1990). Initial ages were therefore drawn from a categorical distribution 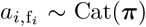 with probabilities that depended on whether age was known. For known ages, *π*_1:10_ ∼ Dir(***α***) with ***α*** = [1, 1, 1, 1, 1, 1, 1, 1, 1, 1]; for unknown ages, *π*_1_ = 0 and *π*_2:10_ ∼ Dir(***α***) with ***α*** = [16, 8, 4, 2, 1, 1, 1, 1, 1]. The latter construct assumes unknown age animals were not first seen in their birth year (given the obvious smaller size of 0.5-year olds) but most likely to be younger than adults. Unknown sex animals had values drawn from the observed sex distribution such that *sex*_*i*_ ∼ Bern(*δ*), where *δ* indicated the probability of being female. Finally, injury status was unknown in years where an individual was not sighted. To estimate the latent injury status, and accommodate some nuance in the probability of injury (Linden et al., 2023), we built a log-linear model:

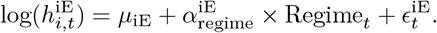

Here, the hazard rate of severe injury by entanglement 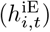 is determined by an intercept (*µ*_iE_), a regime effect for years after 2013 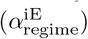, and a random normal effect for year *t* 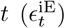. The injury hazard for vessel strikes 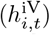 was determined by an intercept only (*µ*_iV_) given the much lower observed rates. The observed rates of severe injury were known to be biased low (Linden et al., 2023), so the corresponding effect sizes of injury on reproduction (*β*_inj_) and survival (*α*_inj_) were expected to be underestimated. As such, we used an informed uniform prior that was restricted to be negative, Uniform (−10, 0), for the effect on calving probability (*β*_inj_) to prevent uncertainty from allowing for unreasonable positive effects (Schofield et al., 2009).

We assigned vague priors for most other parameters, including Gamma(1, 1) for the mortality/injury hazard rate intercepts, Beta(1, 1) for most probabilities, and Normal(0, *σ* = 3) for effects coefficients. We assigned informative priors for the variance parameters of the random error terms where *ϵ* ∼ Normal(0, *σ*); each *σ* was drawn from a scaled Half Student-T (Rankin et al., 2016) with *τ* = 4 and df = 6 (more constrained) for the entanglement injury hazard rates and *τ* = 1 and df = 3 (little constrained) for the mortality hazard rates, the calving probabilities, and the sighting probabilities. The probability of calf loss (*κ*) was assigned a Beta(2, 8) prior to reflect knowledge that it should be low. The probability of a missed summer calf (*ρ*^calf^) was assigned a Beta(5, 445) prior consistent with the observation of 5 calves during summer surveys that were without mothers (individuals that could themselves be sighted without a calf, despite calf survival) out of 450 successful calving events during 1990-2019; this informed prior was a necessary approximation given the lack of an alternative approach to modeling summer calf detection (for additional insights, see Hamilton et al. (2022)). To improve mixing we used a sum-to-zero approach for the random effects (Ogle and Barber, 2020).

The MCMC algorithm was run for 60,000 iterations over 2 chains, after a burn-in of 20,000 iterations. We examined trace plots and the potential scale reduction factor [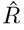; Brooks and Gelman (1998)] to assess convergence and calculated the prior-posterior overlap for the main parameters to assess identifiability (Gimenez et al., 2009). Finally, we assessed model fit with posterior predictive checks (PPC) (Conn et al., 2018) to compare annual counts of events observed to those predicted, using 10,000 posterior samples.

## 3. Results

The 691 individuals across 30 years yielded 7,390 annual detections during summer surveys of whales, with 346 annual detections of females with a calf, across 163 individual adult females. The summer detections of calves represented 96% of known calving events.

Summaries of the posterior distributions for model parameters can be found in Table S3. Trace plots indicated sufficient convergence with all values of 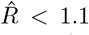. Prior-posterior overlaps were mostly <35% (Table S4) except for the effect of a severe injury on calving (*β*_inj_), and the probability of a missed summer calf (*ρ*^calf^); these two parameters were given informed priors due to sparse information, so the poor identifiability was expected. The posterior predictive check did not indicate any serious lack of fit (Figure S2), though there were several years where observed counts fell outside the envelope of what was predicted. In presenting model results, we report parameter estimates as the posterior median followed with 95% credible intervals in brackets.

### 3.1. Observation processes

Average sighting probability was high with inv-logit(*µ*^*p*^) = 0.792 [0.778, 0.806], consistent with earlier mark-recapture estimates of NARWs on an annual time step (Pace et al., 2017) and with our previous effort at modeling spring/summer surveys only (Linden et al., 2023). Females on average were observed with a lower probability than males 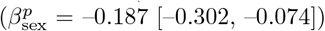 and there was considerable residual variation in the logit-linear model across time (*σ*^*p*[*t*]^ = 0.775 [0.605, 1.048]) and individuals (*σ*^*p*[*i*]^ = 1.171 [1.066, 1.285]). The probability of a missed summer calf was low (*ρ*^calf^ = 0.006 [0.002, 0.013])), and mostly reflected the informed prior distribution as indicated by the high overlap (56%) between the prior and posterior distributions (Table S4).

### 3.2. Variation in injury and survival

The average hazard rate for severe injury by entanglement (exp(*µ*^iE^) = 0.007 [0.005, 0.009]) was much higher than that for vessel strike (exp(*µ*^iV^) = 0.001 [0.000, 0.001]). There was evidence of an elevated entanglement injury rate after 2013 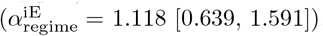.

The average mortality hazard rate for adult males and nulliparous females, exp(*µ*^h^) = 0.010 [0.008, 0.012], translated to an annual survival probability of 0.990 [0.988, 0.992]. The mortality hazard was elevated for younger animals (*α*_1_ = 0.912 [0.400, 1.361]; *α*_2_ = 0.922 [0.404, 1.385]), for females with a calf (*α*_C_ = 0.797 [–0.184, 1.418]), and for females resting from a recent calving event (*α*_R_ = 1.441 [0.983, 1.861]). The presence of a severe injury greatly increased the mortality hazard (*α*_inj_ = 2.980 [2.544, 3.370]), such that annual survival for severely injured animals (e.g., males) was reduced to 0.825 [0.751, 0.884]. Additionally, there was evidence of an elevated mortality rate after 2013 (*α*_regime_ = 0.840 [0.374, 1.263]), even after having accounted for the increased severe injury rate. Combined, an individual with a severe injury after 2013 had an annual survival probability of 0.642 [0.494, 0.767]; prior to 2013, survival of severely injured animals was 0.825 [0.751, 0.884].

### 3.3. Variation in reproduction

The estimated probability of being a female (*δ*) was 0.46 [0.42, 0.50]. The average probability of calving for proven females in a waiting state was inv-logit(*µ*^*B*^) = 0.217 [0.162, 0.281]. There was some evidence that severe injury decreased calving (*β*_inj_ = –1.189 [–4.418, –0.060]), though sparse observations of waiting females with severe injuries motivated the informative prior and resulted in equivocal conclusions. The probability of calving for the first time increased from age 5 (0.001 [0.000, 0.008]) to age 10+ (0.078 [0.051, 0.114]), and there was some evidence of a regime shift in 2010 (*β*_2010regime_ = –0.515 [–1.455, 0.320]) that decreased all stage-specific probabilities of calving (Figure 3). The average annual probability of calving for a proven female decreased from 0.217 [0.162, 0.281] to 0.142 [0.067, 0.252] after the 2010 regime shift. Both prey indices indicated positive relationships with calving, though the GOM prey index had a much larger effect size (*β*_prey=GOM_ = 1.133 [0.521, 1.786]) than the GSL prey index (*β*_prey=GSL_ = 0.307 [–0.079, 0.686]). Additionally, a strong interaction term (*β*_prey=GOM|2010regime_ = –1.149 [–2.168, –0.196]) indicated that the positive relationship for the GOM index disappeared after 2010. Comparing the observed and predicted calving of waiting females illustrated the positive relationships with the each prey index and the change in that relationship in the GOM after 2010 (Figure 4). Finally, calf loss (*κ*) was estimated to be 0.054 [0.039, 0.072].

**Figure 3.**
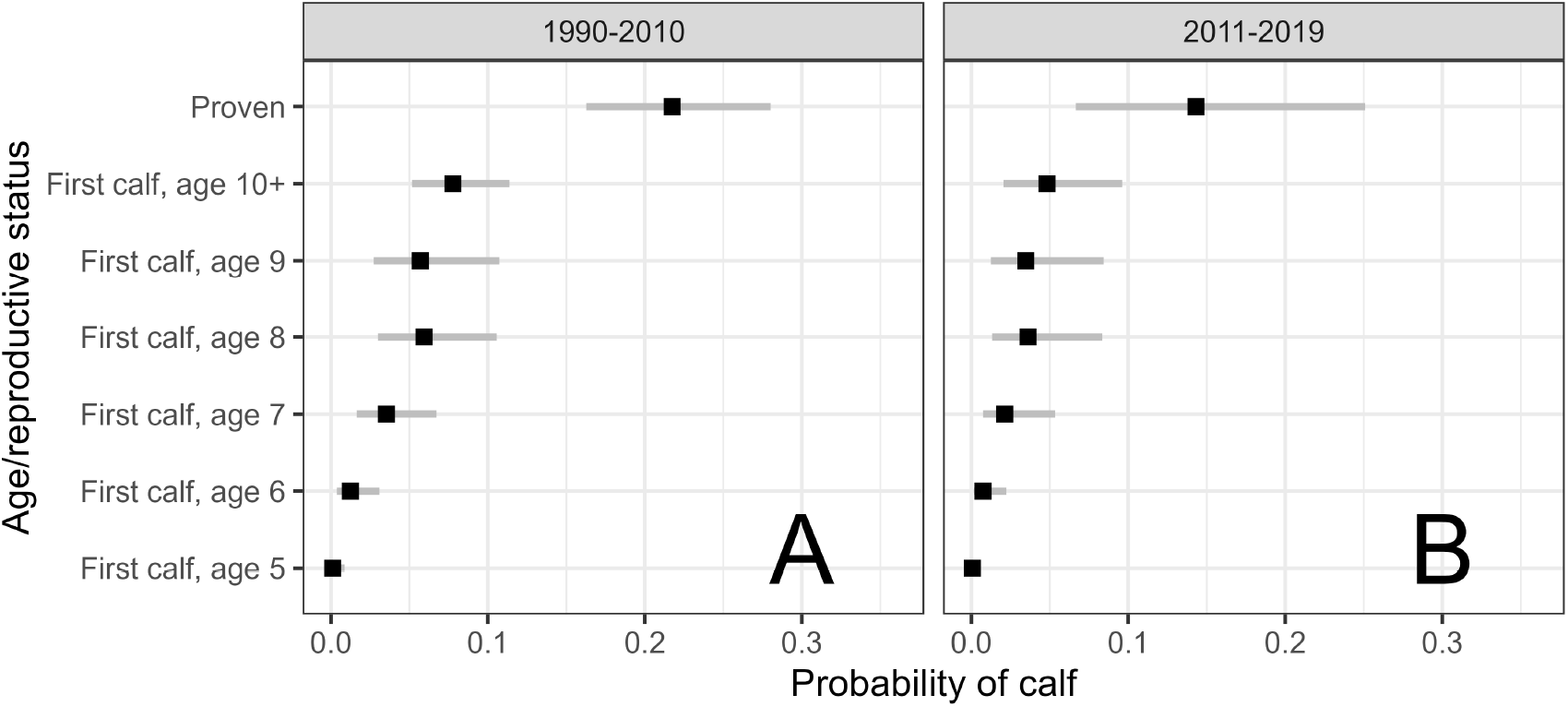
Average probabilities of calving by North Atlantic right whales across age classes for first time breeders (ages 5–10+) and for proven breeders, during average prey availability and for females without severe injuries. Medians with 95% credible intervals presented for 1990-2010 (A) and 2011-2019 (B).

**Figure 4.**
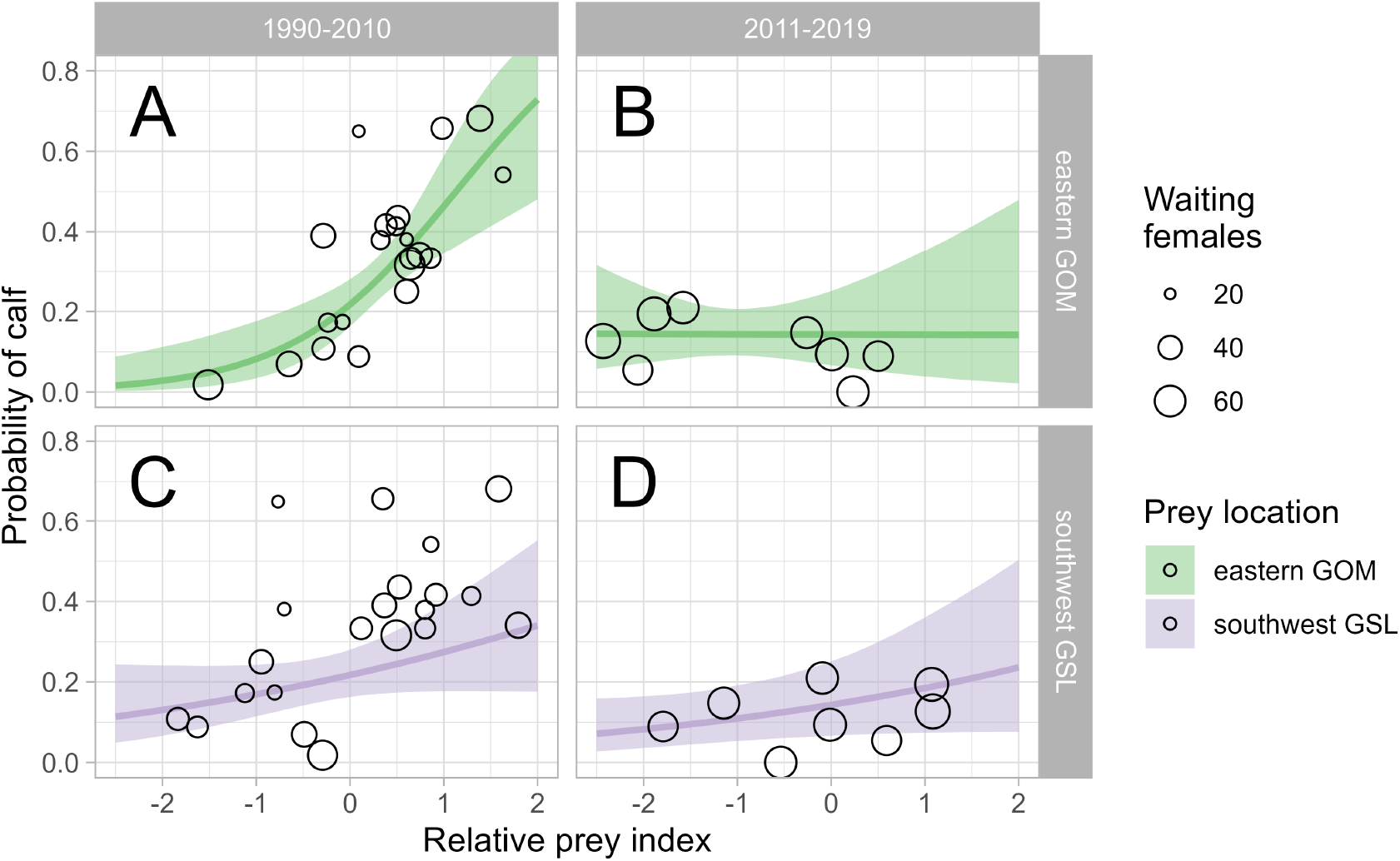
Observed and predicted annual probabilities of calving by North Atlantic right whales across relative prey index values for the eastern Gulf of Maine (A, B) and the southwest Gulf of St. Lawrence (C, D), further separated by temporal regime. Observed values and predictions are for proven females without severe injuries. Median estimate with 95% credible intervals presented for predictions as well as the relative sample size for the proportion of observed waiting females that had a calf.

## 4. Discussion

Our model of North Atlantic right whale reproduction estimated relationships between the probability of calving and multiple individual and temporal attributes relevant to changes in population structure and environmental conditions in the western North Atlantic over the past 30 years. Calving probability had strong positive relationships with indices of *Calanus* spp. biomass in the eastern Gulf of Maine and the southwestern Gulf of St. Lawrence, though the former relationship appears to have broken after 2010. The effect of severe injury on calving was highly uncertain as a result of low sample sizes, in part due to the relatively higher probability of mortality from the injury which would preclude the opportunity for calving. The causes of the recent decline in calving probability (Figure 3) are not fully explained by the factors examined in our model, and therefore the future prospects of the population remain unclear. Population projections suggest both high mortality rates and reduced calving are limiting factors for NARW persistence (Runge et al., 2023).

We made various modeling choices to accommodate a larger effort to represent the population on an annual time step, here in terms of transitions between reproductive states (e.g., sex, with calf, etc.) and elsewhere as a function of cause-specific injury and mortality states (Linden et al., 2023). We chose a “summer” survey period (1 April–30 September) corresponding to when most of the population is observed and used this period to define our encounter histories. We used a 6-month survey period instead of a full “whale year” (Pace et al., 2017) to better accommodate challenges with the closure assumption in capture-recapture modeling (Lebreton et al., 1992), which stipulates no changes in the state of an individual during a survey period. For NARWs, this assumption is most precarious for states of injury and mortality, which in reality are occurring as the result of a continuous hazard through time. Longer sampling periods can improve the quantity of observations and precision in parameter estimates but can also increase violations to the closure assumption, potentially resulting in various biases depending on the ecological process of interest (Dupont et al., 2019). Given an annual reproductive cycle and the extensive survey effort in the NARW breeding area during the boreal winter, there is greater certainty in modeling annual reproductive states at least in terms of females bearing calves (Meyer-Gutbrod et al., 2015; Reed et al., 2022). We used the observed reproductive states during winter surveys as “known states” (Kéry and Schaub, 2012), which improved estimation of calving probability and enabled estimation of calf survival during the first ∼6 months with observational uncertainty. Still, the inability to demonstrate a clear relationship between severe injury and calving was likely a result of trade-offs made in defining encounter histories for the analysis.

Other modeling efforts have used smaller survey periods (e.g., 3 months) to model fine-scale transitions between health states and estimate the role of injuries in NARW demographic rates (Schick et al., 2013; Knowlton et al., 2022; Pirotta et al., 2023). The role of severe vs. minor/moderate injuries and the difficulty in estimating the effects of the former on calving was well illustrated by Pirotta et al. (2023): severe injuries that lead to death cannot be used to assess calving. Our model did not consider injuries classified as less than severe in the probability of calving, which may be a priority for further development. Severe injury observations are an underestimate of true injury rates (Linden et al., 2023), and the same is likely true of minor and moderate injuries. Assuming similar patterns in rates of injury over time regardless of severity, the observed increase in severe injuries since 2013 could correspond to an increase in minor/moderate injuries and partially explain the negative regime effect on calving probability.

Additional explanations for the 2010 regime effect could include a long-term trend in declining reproduction due to challenges related to the quality and quantity of prey not explained by our prey indices. The binary before vs. after indicator for regime would result in a similar regression coefficient to a linear trend, at least in terms of sign. While no long-term trend was apparent (Figure 2), other empirical evidence suggests a reduction in the nutritional value of *Calanus* spp. (McKinstry et al., 2013; Helenius et al., 2024). Reduced blubber thickness in comparison to southern right whales (Miller et al., 2011) and long-term decreases in body length (Stewart et al., 2021) suggest the potential for large-scale nutritional deficiencies that may be temporarily overcome during periods of high prey availability (see Figure 4 in Pirotta et al., 2023), but ultimately contribute to longer calving intervals and reduced reproduction. Reed et al. (2022) attributed the recent pattern of decreased calving probability to an increasing failure to start breeding by females born after 2000; the authors hypothesized potential mechanisms related to the decreasing size of whales and higher entanglement rates (particularly of breeding females). We should note that our modeling of calf loss did not consider perinatal mortality, which may also explain increases in delayed first parturition (females having their first calf) and calving intervals (Browning et al., 2010).

It is very likely that our chosen prey indices provide an incomplete picture of prey availability, given the complex spatial variability in NARW habitat and zooplankton community dynamics that are likely related to NARW resource acquisition (Meyer-Gutbrod et al., 2015, 2023). There are many challenges with modeling climate-mediated habitat conditions that vary over large spatial and temporal scales and have myriad forms of potential summarization, some of which can be overcome using an explicit predictive framework (Gerber et al., 2015). The strong positive relationship between the GOM index and calving is unsurprising, given the historical importance of the GOM for summer feeding (Meyer-Gutbrod et al., 2023). Meyer-Gutbrod et al. (2015) previously demonstrated a strong association between NARW calving probability and *Calanus finmarchicus* abundance in the GOM through 2007 while accommodating more complexity and nuance in the seasonal and spatial variation of zooplankton dynamics. Our estimated change in this relationship after 2010 follows logically from the large ecosystem shift in the western Atlantic that likely resulted in a distribution shift by NARW shortly after (Sorochan et al., 2019). Still, it is unclear why the prey index value for the summer following the calving season would have explanatory power; it may indicate some association with large scale environmental conditions over multiple years that are favorable to resource acquisition, or possibly suggest that right whales are capable of anticipating a pulse in food resources (Bergeron et al., 2011). The evidence of a positive calving relationship with the GSL index may seem curious given that regular use of this region by NARWs is a relatively new phenomenon (Meyer-Gutbrod et al., 2023); the *Calanus* spp. biomass in the GSL likely serves as coarse proxy of the true mechanisms (e.g., Greene et al., 2008), or an indicator of eventual conditions downstream from the Scotian shelf (e.g., in the GOM).

The prospect for NARW recovery is currently constrained by anthropogenic mortality (Corkeron et al., 2018), but it remains unclear how much the more recent unexplained decline in reproduction is due to human-caused interactions (e.g., chronic sub-lethal entanglement) versus large-scale trends in habitat quality resulting from climate change (Meyer-Gutbrod et al., 2021). Our reproduction model provides demographic parameter estimates that can be used in population projections, though uncertainty in the mechanisms driving the post-2010 regime effect will limit interpretations of future population trajectories. For example, if the observed stunting in individual growth is irreversible (Stewart et al., 2021), the regime shift would represent a new reality for the species that will constrain, and may decrease, population growth in the future. Entanglement mitigation could improve mortality rates in the short term, but the effects on stunting and delayed first parturition would likely be postponed at best. Apparent nutritional deficiencies resulting from climate change (Miller et al., 2011; Gavrilchuk et al., 2021) are unlikely to improve, though NARWs may be exhibiting some resilience to change (Bishop et al., 2022). The need for an ecosystem-based management perspective remains crucial to setting realistic NARW recovery goals (Meyer-Gutbrod et al., 2015).

## Acknowledgements

We are grateful to the North Atlantic Right Whale Consortium (NARWC) for access to the sightings data. The capacity to develop precise estimates of North Atlantic right whale demographic parameters is due to the thousands of photographic captures of whales contributed by hundreds of collaborators working through the NARWC for nearly 40 years. We thank Keven Sorochan and Stéphane Plourde for sharing their knowledge on *Calanus* ecology and future trends, and for kindly processing and providing both published and unpublished *Calanus* data. We would also like to acknowledge the following data curators of the AZMP (DFO) swGSL-June mackerel egg survey and EcoMon (NOAA) who kindly provided the *Calanus* data: David Bélanger (DFO, NL), Benoit Casault (DFO, NL), Caroline Lehoux (DFO, Qc), Harvey Walsh (NOAA). We thank reviewers from the Center for Independent Experts for helpful comments on an earlier version of this work. Any use of trade, firm, or product names is for descriptive purposes only and does not imply endorsement by the U.S. Government.

## Supplementary Material

*Disclaimer: Any use of trade, firm, or product names is for descriptive purposes only and does not imply endorsement by the U.S. Government*.

**Figure S1:**
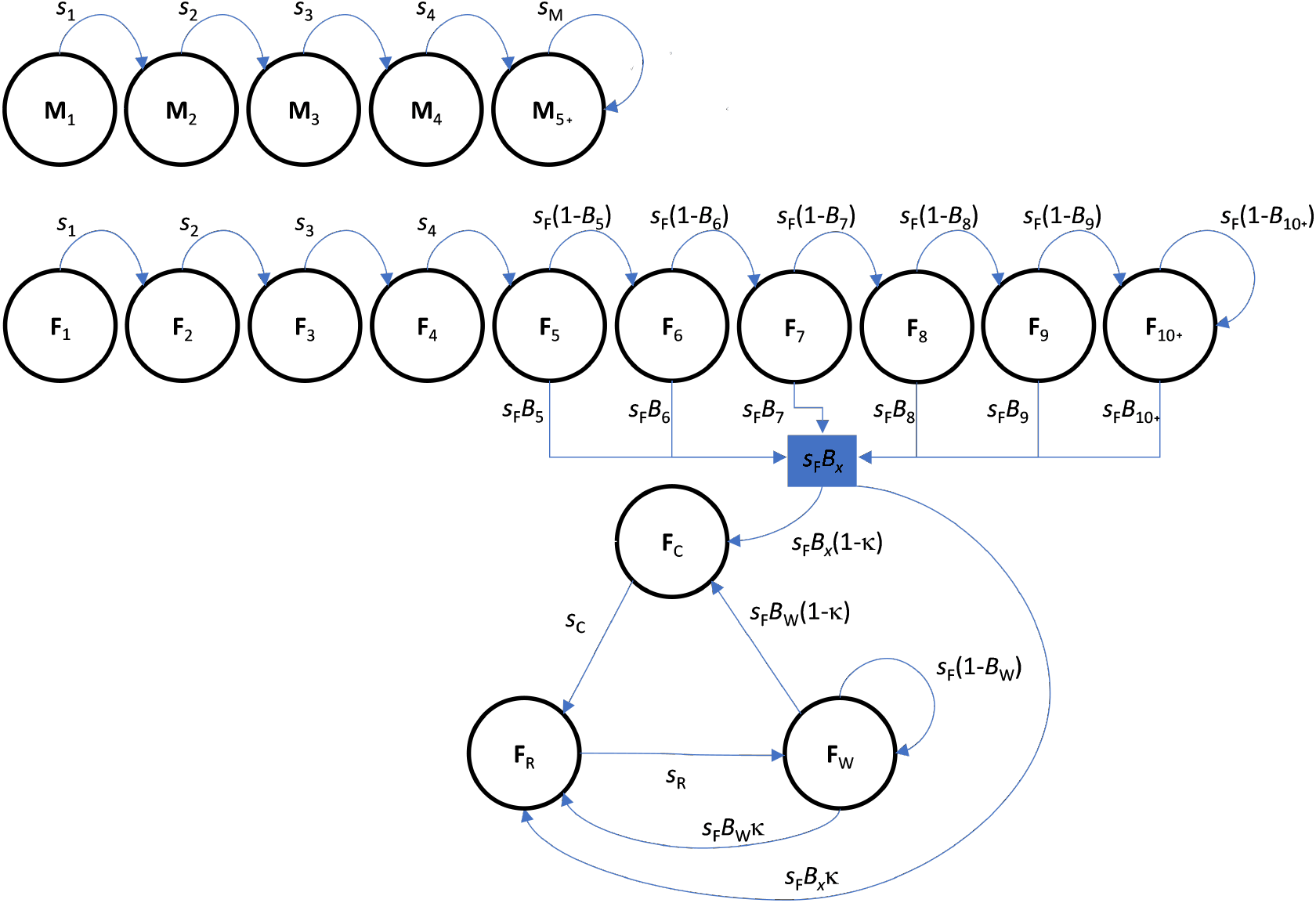
State transition model for North Atlantic right whale reproductive states and age classes across annual time steps. Transitions between states from year *t* to *t* + 1 depend on probabilities of survival (*s*) and reproduction (*B*), which are conditional on the state and age class of an individual in year *t*. Transitions to mortality and subscripts for indexing individual *i* and year *t* are omitted for clarity.

## Appendix S1: Exploration of relationships between *Calanus* indices and per-capita calving

Here, we explored several regional *Calanus* spp. indices and their ability to explain variation in per-capita probability that an adult female would successfully birth a calf.

## 0.0.1. Calanus *data*

The data included predictions from Sorochan et al. (2019) of *Calanus* spp. biomass at 4 subregions: south-west Gulf of St. Lawrence (swGSL), eastern Gulf of Maine (eGOM), western Gulf of Maine (wGOM), and Georges Bank (GOMGB). Sorochan et al. (2019) describe the monitoring programs that collected the zooplankton samples, including the Fisheries and Oceans Canada mackerel egg production survey in the swGSL and the National Oceanic and Atmospheric Administration Marine Resources Monitoring, Assessment and Prediction and Ecosystem Monitoring surveys (“EcoMon”) in the eGOM, wGOM, and GOMGB. Biomass (mg m^-2^) of the fourth and fifth copepodite and adult stages was derived from stage-specific abundance (individuals m^-2^) and individual dry weight (mg individual^-1^). Annual estimates were calculated from general linear models to combine values across seasons (and stations) for each subregion. The published data through 2016 in Sorochan et al. (2019) were kindly supplemented through 2019 for the the purposes of our modeling (K. Sorochan, Fisheries and Oceans Canada, *unpublished data*, 2022). The annual values for each subregion were total *Calanus* spp. biomass on the log scale.

## 0.0.2. Right whale calving

Observed calf counts of North Atlantic right whales are documented each boreal winter season during extensive aerial and boat surveys by the North Atlantic Right Whale Consortium (https://www.narwc.org/) and its various partners and contributors (Kraus and Rolland, 2007). We combined these counts with an annual estimate of the number of adult NARW females during 1990–2019 generated from the mark-recapture model used for population assessment (Pace et al., 2017; Linden, 2023). While not all adult females (age 5+) are capabable of breeding, we considered the count of adult females to be a reasonable proxy.

The following table illustrates the data used for exploration:

| year | Right whales |  | Calanus index (log biomass) |  |  |  |
| --- | --- | --- | --- | --- | --- | --- |
|  | calves | NF | GOMeast | GOMGB | GOMwest | swGSL |
| 1985 | 11 | - | 2.961 | 2.190 | 2.938 | 2.617 |
| 1986 | 13 | - | 3.000 | 1.912 | 3.090 | 2.981 |
| 1987 | 11 | - | 3.194 | 2.435 | 3.027 | 2.192 |
| 1988 | 7 | - | 3.186 | 2.539 | 2.747 | 3.141 |
| 1989 | 16 | - | - | 2.212 | 2.559 | 2.408 |
| 1990 | 12 | 92 | - | 2.185 | 2.429 | 2.652 |
| 1991 | 17 | 95 | 3.140 | 2.108 | 2.681 | 2.909 |
| 1992 | 12 | 104 | 3.240 | 2.359 | 2.845 | 2.500 |
| 1993 | 6 | 101 | 2.970 | 2.201 | 2.705 | 2.530 |
| 1994 | 9 | 112 | 3.190 | 2.208 | 2.923 | 1.965 |
| 1995 | 7 | 117 | 3.134 | 2.259 | 3.013 | - |
| 1996 | 21 | 119 | 3.246 | 2.354 | 2.945 | 3.002 |
| 1997 | 19 | 118 | 3.204 | 2.285 | 3.010 | - |
| 1998 | 5 | 118 | 2.819 | 2.398 | 2.757 | 1.695 |
| 1999 | 4 | 116 | 3.134 | 2.085 | 2.965 | 3.287 |
| 2000 | 1 | 114 | 2.960 | 2.297 | 2.904 | 2.980 |
| 2001 | 31 | 123 | 3.432 | 2.349 | 3.392 | 3.124 |
| 2002 | 21 | 127 | 3.093 | 2.286 | 3.355 | 3.319 |
| 2003 | 19 | 126 | 3.328 | 2.562 | 3.360 | 3.362 |
| 2004 | 16 | 123 | 3.215 | 2.721 | 3.327 | 3.044 |
| 2005 | 28 | 124 | 3.135 | 2.470 | 3.047 | 2.763 |
| 2006 | 19 | 127 | 3.138 | 2.154 | 3.036 | 3.703 |
| 2007 | 23 | 135 | 3.261 | 2.394 | 3.262 | 3.593 |
| 2008 | 23 | 134 | 3.212 | 2.115 | 3.194 | 3.541 |
| 2009 | 39 | 142 | 3.332 | 2.205 | 3.037 | 3.477 |
| 2010 | 19 | 149 | 2.920 | 2.230 | 3.042 | 2.634 |
| 2011 | 22 | 151 | 3.006 | 2.252 | 2.959 | 2.985 |
| 2012 | 7 | 150 | 2.839 | 1.895 | 2.964 | 3.638 |
| 2013 | 20 | 164 | 2.972 | 2.092 | 2.984 | 3.322 |
| 2014 | 11 | 172 | 2.844 | 2.341 | 2.876 | 3.003 |
| 2015 | 17 | 172 | 3.087 | 2.046 | 2.904 | 2.075 |
| 2016 | 14 | 165 | 3.325 | 2.135 | 3.470 | 2.291 |
| 2017 | 5 | 158 | 3.114 | 2.259 | 2.933 | 2.497 |
| 2018 | 0 | 145 | 2.995 | 2.319 | 3.318 | 3.236 |
| 2019 | 7 | 138 | 3.267 | 2.087 | 3.163 | 2.993 |

## 0.0.3. Binomial regression modeling

We used binomial regressions of calves per female to identify a useful covariate structure for the multistate capture-recapture model. Our multi-stage process included identifying the size and potential shift in the multi-year window of *Calanus* index values that could best predict calving probability. Once we identified the ideal window, we further refined the sub-region selection and explored an interaction with temporal regime (pre/post 2010).

The probability mass function for a binomial random variable, *Y*, is defined as:

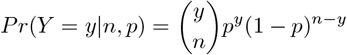

where *y* is the number of successes (e.g., calves born), *n* is the number of trials (e.g., adult females), and *p* is the probability of success. Variation in *p* can be accommodated by a logit-linear model:

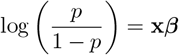

where **x** is a collection of predictor variables and ***β*** is a vector of coefficients describing the relationships between the predictors and probability of success. We fit the binomial regressions in R (R Core Team, 2022) and used calculations of AIC and proportion of deviance explained to compare models. The general form of the models in R was as follows:

~~~
glm(cbind(calves,NF-calves) ∼ GOMeast + GOMGB + GOMwest + swGSL,
    family = binomial, weights = NF)
~~~

To identify the multi-year window shift and size, we calculated rolling averages of *Calanus* values across window sizes of 1 to 5, with values shifted 0 to 2 years. The rolling average involved exponentiating the index values, taking the mean, and then log-transforming the result. The following table displays the model performance across each window shift and size:

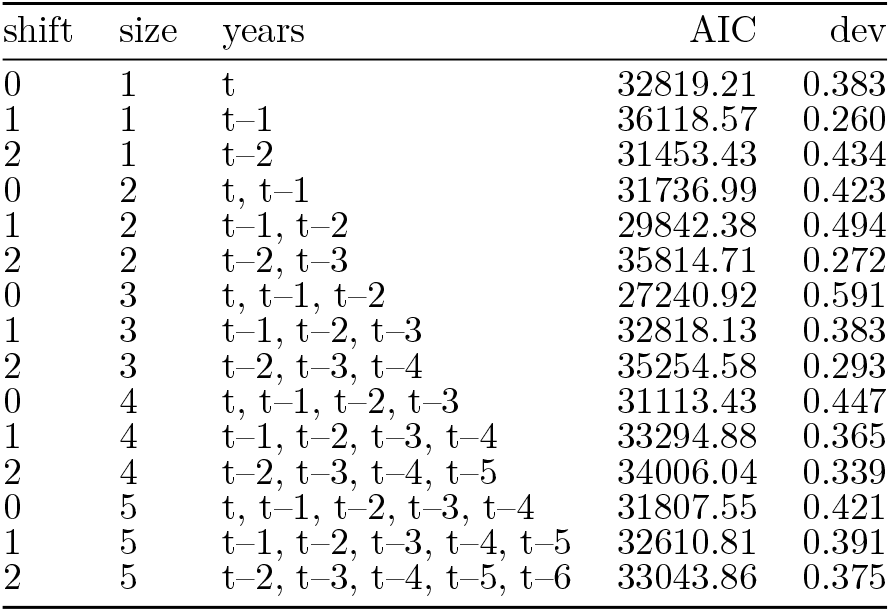

| shift | size | years | AIC | dev |
| --- | --- | --- | --- | --- |
| 0 | 1 | t | 32819.21 | 0.383 |
| 1 | 1 | t-1 | 36118.57 | 0.260 |
| 2 | 1 | t-2 | 31453.43 | 0.434 |
| 0 | 2 | t, t-1 | 31736.99 | 0.423 |
| 1 | 2 | t-1, t-2 | 29842.38 | 0.494 |
| 2 | 2 | t-2, t-3 | 35814.71 | 0.272 |
| 0 | 3 | t, t-1, t-2 | 27240.92 | 0.591 |
| 1 | 3 | t-1, t-2, t-3 | 32818.13 | 0.383 |
| 2 | 3 | t-2, t-3, t-4 | 35254.58 | 0.293 |
| 0 | 4 | t, t-1, t-2, t-3 | 31113.43 | 0.447 |
| 1 | 4 | t-1, t-2, t-3, t-4 | 33294.88 | 0.365 |
| 2 | 4 | t-2, t-3, t-4, t-5 | 34006.04 | 0.339 |
| 0 | 5 | t, t-1, t-2, t-3, t-4 | 31807.55 | 0.421 |
| 1 | 5 | t-1, t-2, t-3, t-4, t-5 | 32610.81 | 0.391 |
| 2 | 5 | t-2, t-3, t-4, t-5, t-6 | 33043.86 | 0.375 |

Results suggested that a 3-yr window with no shift (i.e., including the focal year, *t*, and previous 2 years) provided the best model fit in terms of AIC and proportion of deviance explained. Parameter estimates for that model were as follows:

~~~
summary(mod1)
Call:
glm(formula = cbind(calves, NF - calves) ∼ scale(GOMeast) + scale(GOMGB) +
 scale(GOMwest) + scale(swGSL), family = binomial, data = dat.glm,
 weights = NF)
Coefficients:
 Estimate Std. Error z value Pr(>|z|)
(Intercept) −2.104497 0.004734 −444.56 <2e-16 ***
scale(GOMeast) 0.344039 0.005408 63.61 <2e-16 ***
scale(GOMGB) 0.097399 0.005556 17.53 <2e-16 ***
scale(GOMwest) −0.322273 0.006645 −48.50 <2e-16 ***
scale(swGSL) 0.471756 0.005425 86.95 <2e-16 ***
---
Signif. codes: 0 ‘***’ 0.001 ‘**’ 0.01 ‘*’ 0.05 ‘.’ 0.1 ‘ ‘ 1
(Dispersion parameter for binomial family taken to be 1)
 Null deviance: 26883 on 29 degrees of freedom
Residual deviance: 11006 on 25 degrees of freedom
AIC: 27241
Number of Fisher Scoring iterations: 5
~~~

We decided to remove both the GOMGB and GOMwest indices; the former due to its relatively small effect size, and the latter due to its unexpected negative relationship. Including an interaction with the 2010 regime shift yielded a final model:

~~~
summary(mod2)
Call:
glm(formula = cbind(calves, NF - calves) ∼ scale(GOMeast) * regime2010 +
 scale(swGSL), family = binomial, data = dat.glm, weights = NF)
Coefficients:
 Estimate Std. Error z value Pr(>|z|)
(Intercept) −2.177679 0.007852 −277.34 <2e-16 ***
scale(GOMeast) 0.600824 0.011006 54.59 <2e-16 ***
regime2010 −0.443711 0.014728 −30.13 <2e-16 ***
scale(swGSL) 0.144578 0.006426 22.50 <2e-16 ***
scale(GOMeast):regime2010 −0.693226 0.016412 −42.24 <2e-16 ***
---
Signif. codes: 0 ‘***’ 0.001 ‘**’ 0.01 ‘*’ 0.05 ‘.’ 0.1 ‘ ‘ 1
(Dispersion parameter for binomial family taken to be 1)
 Null deviance: 26883 on 29 degrees of freedom
Residual deviance: 10996 on 25 degrees of freedom
AIC: 27231
Number of Fisher Scoring iterations: 5
~~~

## 0.0.4. ConclusionsExploration of relationships between

This exploratory analysis suggested that probability of calving had a positive relationship with *Calanus* indices in the swGSL and GOMeast. After 2010, the average probability of calving decreased and the GOMeast index no longer explained variation. While the exploration was informative, it does not consider whether the adult females considered as “trials” in the binomial regression were actually available to breed (e.g., if an individual has a calf in year *t*, it cannot have a calf in year *t* + 1). Still, the covariate structure served as a guide for the multistate capture-recapture model where calving availability is explicitly addressed.

**Table S3:** Parameter estimates from a multistate capture-recapture model of North Atlantic right whale survival and reproduction during 1990–2019. Posterior summaries include mean, standard deviation (SD), and quantiles describing the median and lower/upper 95% intervals.

| parameter | mean | SD | 2.5% | 50% | 97.5% |
| --- | --- | --- | --- | --- | --- |
| mu.h | -4.631 | 0.110 | -4.854 | -4.629 | -4.426 |
| mu.iE | -5.018 | 0.166 | -5.353 | -5.014 | -4.709 |
| mu.iV | -7.376 | 0.474 | -8.405 | -7.338 | -6.556 |
| alpha.age[1] | 0.905 | 0.244 | 0.400 | 0.912 | 1.361 |
| alpha.age[2] | 0.914 | 0.248 | 0.404 | 0.922 | 1.385 |
| alpha.age[3] | -0.344 | 0.429 | -1.293 | -0.314 | 0.418 |
| alpha.age[4] | 0.144 | 0.720 | -1.594 | 0.285 | 1.007 |
| alpha.C | 0.751 | 0.417 | -0.184 | 0.797 | 1.418 |
| alpha.R | 1.437 | 0.224 | 0.983 | 1.441 | 1.861 |
| alpha.inj | 2.976 | 0.209 | 2.544 | 2.980 | 3.370 |
| alpha.2013regime | 0.835 | 0.224 | 0.374 | 0.840 | 1.263 |
| alpha.iE.2013regime | 1.117 | 0.243 | 0.639 | 1.118 | 1.591 |
| mu.B | -1.284 | 0.179 | -1.645 | -1.282 | -0.940 |
| beta.age.5 | -5.712 | 1.423 | -9.041 | -5.502 | -3.528 |
| beta.age.6 | -3.124 | 0.519 | -4.237 | -3.089 | -2.200 |
| beta.age.7 | -2.041 | 0.347 | -2.759 | -2.026 | -1.404 |
| beta.age.8 | -1.489 | 0.307 | -2.120 | -1.479 | -0.913 |
| beta.age.9 | -1.529 | 0.343 | -2.247 | -1.514 | -0.888 |
| beta.age.10 | -1.194 | 0.163 | -1.518 | -1.193 | -0.880 |
| beta.inj | -1.459 | 1.169 | -4.418 | -1.189 | -0.060 |
| beta.prey=GSL | 0.305 | 0.194 | -0.079 | 0.307 | 0.686 |
| beta.prey=GOM | 1.140 | 0.320 | 0.521 | 1.133 | 1.786 |
| beta.2010regime | -0.529 | 0.448 | -1.455 | -0.515 | 0.320 |
| beta.prey=GOM\|2010regime | -1.158 | 0.497 | -2.168 | -1.149 | -0.196 |
| kappa | 0.054 | 0.008 | 0.039 | 0.054 | 0.072 |
| mu.p | 1.337 | 0.042 | 1.255 | 1.337 | 1.421 |
| beta.p.sex | -0.188 | 0.058 | -0.302 | -0.187 | -0.074 |
| rho.calf | 0.007 | 0.003 | 0.002 | 0.006 | 0.013 |
| delta | 0.459 | 0.019 | 0.422 | 0.459 | 0.496 |
| sigma.iE.t | 0.123 | 0.097 | 0.013 | 0.097 | 0.374 |
| sigma.h.t | 0.330 | 0.102 | 0.148 | 0.324 | 0.554 |
| sigma.B.t | 0.660 | 0.156 | 0.402 | 0.645 | 1.008 |
| sigma.p.t | 0.789 | 0.113 | 0.605 | 0.775 | 1.048 |
| sigma.p.ind | 1.172 | 0.056 | 1.066 | 1.171 | 1.285 |

**Table S4:**
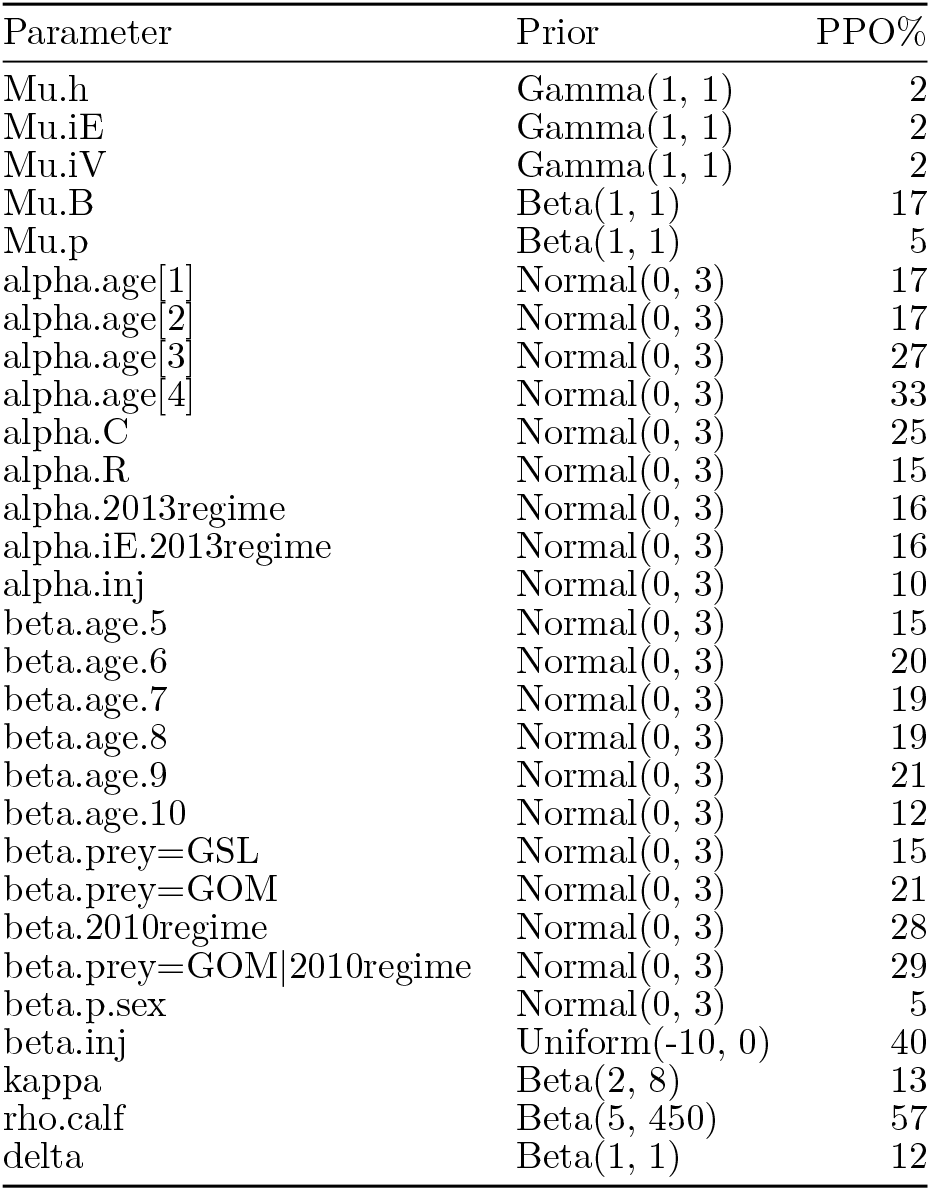
Percentages of prior-posterior overlap for main parameters. While *µ* values were link-scale intercepts, all priors were specified on the real scale of the parameter (e.g., annual probability or rate) and required transformation.

**Figure S2:**
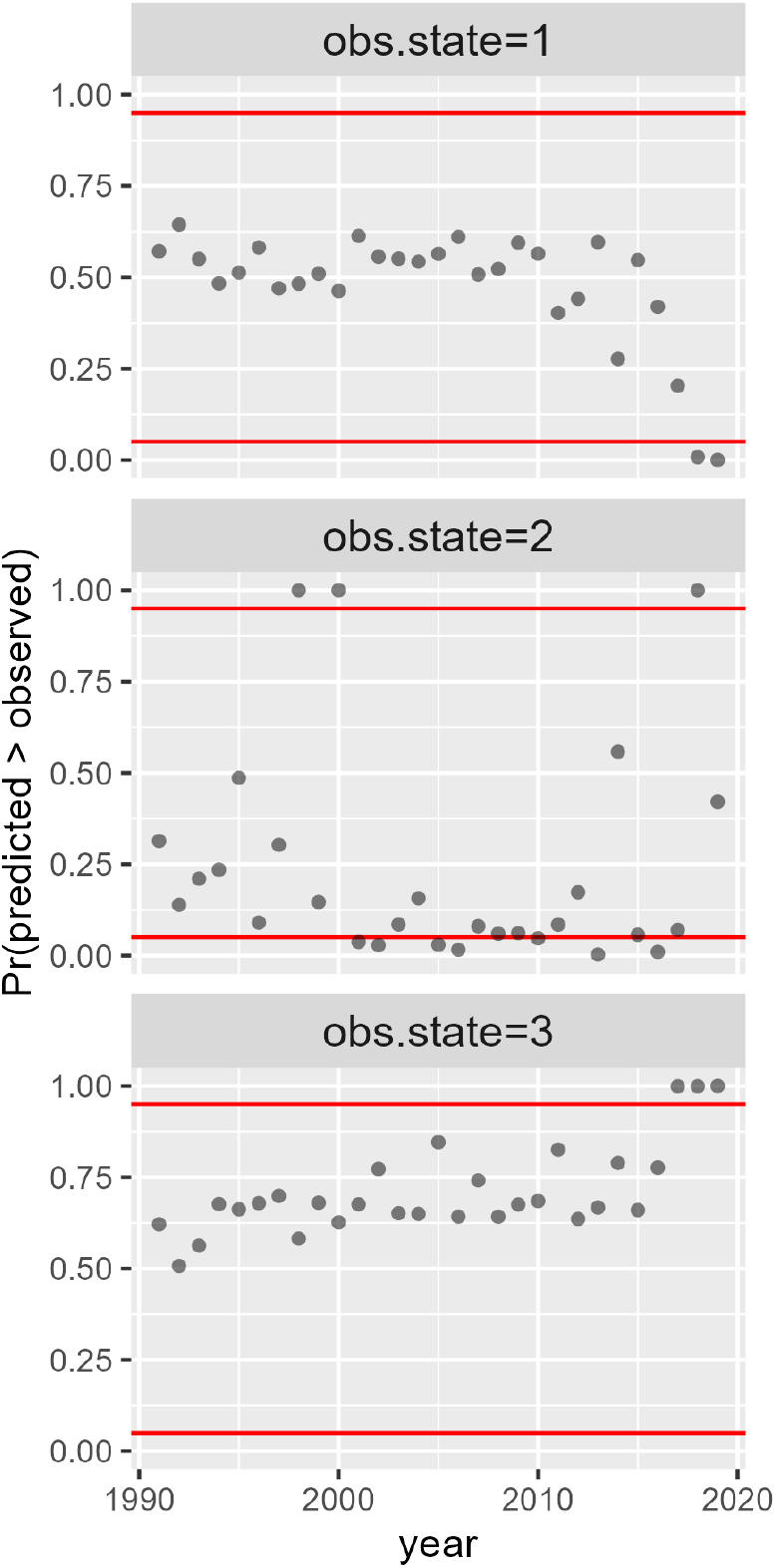
Posterior predictive check illustrating the proportion of simulated data sets that had a predicted count of events greater than the observed count of events for each event type (e.g., observation state) and year. Red lines indicate upper and lower thresholds (0.95 and 0.05, respectively) between which goodness-of-fit is adequate. Recorded events include: 1 = seen alive; and 2 = seen alive with calf.

